# Guiding biomolecular interactions in cells using *de novo* protein-protein interfaces

**DOI:** 10.1101/486902

**Authors:** Abigail J. Smith, Franziska Thomas, Deborah Shoemark, Derek N. Woolfson, Nigel J. Savery

## Abstract

An improved ability to direct and control biomolecular interactions in living cells would impact on synthetic biology. A key issue is the need to introduce interacting components that act orthogonally to endogenous proteomes and interactomes. Here we show that low-complexity, *de novo* designed protein-protein-interaction (PPI) domains can substitute for natural PPIs and guide engineered protein-DNA interactions in *Escherichia coli.* Specifically, we use *de novo* homo- and hetero-dimeric coiled coils to reconstitute a cytoplasmic split adenylate cyclase; to recruit RNA polymerase to a promoter and activate gene expression; and to oligomerize both natural and designed DNA-binding domains to repress transcription. Moreover, the stabilities of the heterodimeric coiled coils can be modulated by rational design and, thus, adjust the levels of gene activation and repression *in vivo.* These experiments demonstrate the possibilities for using designed proteins and interactions to control biomolecular systems such as enzyme cascades and circuits in cells.

## INTRODUCTION

The advent of synthetic biology has brought an increased demand for protein components of reduced size and complexity, which are orthogonal to cellular systems and that function according to understood parameters. Protein-protein interactions (PPIs) are one aspect of protein function that is amenable to design and manipulation. Moreover, an ability to design PPIs completely *de novo* and predictably would impact broadly in synthetic biology by allowing biomolecular interactions and functions to be guided and orchestrated in cells with precision and, potentially, without interfering with endogenous proteomes and interactomes. Whilst excellent progress has been made on the *de novo* design and assembly of PPI-mediated macromolecular structures *in vitro*,^1–4^ much less has been done in living cells. Success here would allow the targeting of proteins to prescribed cellular regions, the co-localization of enzymes to optimize bioproduction, the reconstitution of split proteins to switch enzyme activity on and off, and the assembly of completely new structures in cells to act as scaffolds or compartments for such processes.^5–8^ An advantage of targeting PPIs to take control in synthetic biology is that the PPI components are usually separable from the downstream activity, and so designed PPIs will find applications across many different systems.

An important example of PPIs in cells is transcription control, where PPI-mediated recruitment of components underlies most forms of gene activation.^9^ Transcription repression is also often underpinned by PPIs, either by recruitment of corepressors or because the multimerization of the repressor proteins is a prerequisite for DNA binding.^10,11^ Indeed, in cell and synthetic biology, transcription regulation has provided proof-of-concept systems in which to monitor and exploit PPIs within cells.^12–15^ In their simplest forms, transcription activators consist of a DNA-binding domain, which defines the promoter-specificity of its action, and a PPI domain that recruits RNA polymerase (RNAP) or an associated factor.^14^ Bacterial repressor proteins are conceptually even simpler, as an isolated DNA-binding domain can repress transcription by sterically blocking RNAP binding. However, most natural bacterial repressor proteins exist as PPI-dependent multimers. The cooperative binding that results from multimerization can be important for the design and function of Gene Regulatory Networks (GRNs).^16^ For both activators and repressors, the affinity of the PPI and of the protein-DNA interaction are key parameters that define the behavior of the components within such GRNs.

One of the best-understood PPI motifs is the α-helical coiled coil (CC).^17,18^ This understanding has led to considerable success in CC design.^4,19,20^ CCs are abundant in nature and usually display heptad sequence repeats of hydrophobic (h) and polar (p) amino acids, *hpphppp* (often denoted *abcdefg).* These repeating patterns direct the folding of amphipathic α helices, which assemble *via* their hydrophobic faces to form left-handed rope-like structures with two or more helices in parallel or antiparallel orientations.^17,18^ The rules that govern assembly of CCs have been deciphered.^19–21^ In turn, these have enabled the rational design of “toolkits” of CC peptides that assemble in homo- or hetero-multimeric complexes predictably *in vitro^22–27^* In one such study from one of our laboratories, the hydrophobic amino acids at positions *a* and *d* have been varied to create a set of 30-residue peptides that form parallel homomeric dimers, trimers and tetramers, which have been characterized to atomic resolution.^23^ These peptides are named CC-Di, CC-Tri and CC-Tet, respectively. A series of parallel heterodimeric CCs has also been designed, in which one set of peptides has acidic amino acids at the *e* and *g* positions and another complementary set has basic residues at the *e* and *g* sites.^24^ These CC-Di-A and CC-Di-B peptides do not fold in isolation, but combine when mixed to form stable, obligate heterodimers. Moreover, as CC stability increases with increasing chain length, the CC-Di-AB heterodimers can be tuned to give a range of dissociation constants that varies over several orders of magnitudes *in vitro*^24^

Natural and synthetic CCs have been shown to function effectively as PPIs within transcription activators in yeast and *E. coli.*^25,28^ Here, we test the ability of the *de novo* designed homo and hetero-dimeric CC peptides to function as PPI domains in a range of contexts within living *E. coli* cells. We find that the CC peptides can mediate PPIs in multiple systems *in vivo*, including as part of a cytoplasmic split enzyme, and as components of both transcription repressors and transcription activators. In most cases, the binding affinity designed and measured *in vitro* is reflected in the strengths of the regulatory activity measured *in vivo.* The heterodimeric sequences show the expected specificity, with little or no self-association or off-target activity evident. To demonstrate the complete modular design of synthetic transcription factors, we combine the *de novo* CC-based PPIs with programmable DNA-binding domains based on TAL repeats to generate homo- and heterodimeric transcription regulators. Thus, in these artificial transcription factors, both PPI activity and DNA-binding activity can be designed to match the requirements of a desired application.

## RESULTS AND DISCUSSION

### Protein Colocalization *In Vivo* by *De Novo* Designed PPIs

First, we measured the ability of our toolkit of CC peptides to bring together the components of a split cytoplasmic enzyme. In this system the adenylate cyclase protein of *Bordetella pertussis* is expressed as two separate domains (T25 and T18), which come together to form an active enzyme when fused to partner proteins that form a PPI.^29^ This reconstitution produces cyclic AMP (cAMP), which is detected by monitoring the production of a *lacZ* reporter gene regulated by the cAMP receptor protein (CRP) (Figure 1a).

**Figure 1.**
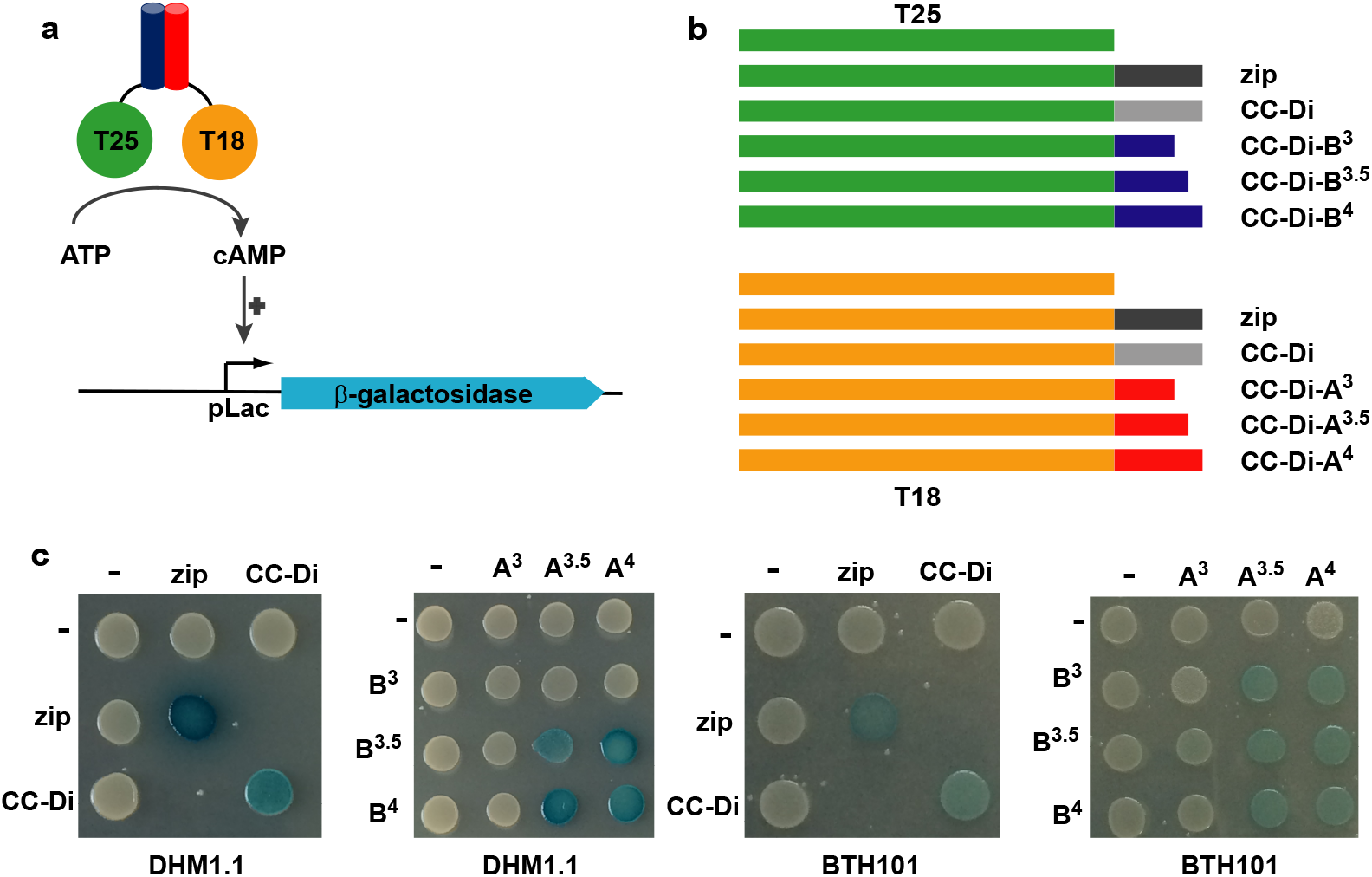
*De novo* designed PPIs interact *in vivo.* (a) The T25 and T18 domains of *B. pertussis* adenylate cyclase can be reconstituted in the presence of interacting CC peptides and this positively regulates expression of β-galactosidase. (b) Fusion proteins used in this assay. The T25 or T18 subdomains are fused to CCs via a short linker. CCs are labelled as follows: Zip, leucine zipper of the yeast GCN4 protein; CC-Di, homodimeric coiled coil; CC-Di-A^x^ and CC-Di-B^x^, acidic or basic heterodimeric CCs comprising x heptad repeats. (C) The *cya^-^ E. coli* stains DHM1.1 and BTH101 were transformed with pairs of fusion proteins as indicated and cultures were spotted on LB agar containing X-gal + IPTG. Blue coloration indicates production of β-galactosidase. The horizontal labels indicate T18-CC-Di-A fusions and the vertical labels indicate T25-CC-Di-B fusions.

We tested two *de novo* CC PPIs: the homodimeric CC-Di^23^ and the heterodimer CC-Di-AB system, in which complementary CC-Di-A and CC-Di-B peptides have been made with 3-, 3.5- and 4-heptad repeats.^24^ Plasmids encoding these CC peptides fused to the *C* termini of the components of split adenylate cyclase *(i.e.,* the T25 and T18 domains) were constructed using sequences codon-optimized for expression in *E. coli* (Figure 1b and Table S1). Reconstitution of adenylate cyclase was monitored in an *E. coli* strain DMH1.1, which lacks the native adenylate cyclase gene (cya^-^). T25 and T18 fusions were co-expressed, and expression of β-galactosidase from the cAMP-dependent *lacZ* gene was monitored by the production of a blue colony phenotype when the transformants were grown on rich X-gal indicator agar.

Cells expressing T25 and T18 without fusion partners did not produce detectable β-galactosidase (Figure 1c), and a positive control with these components fused to the yeast GCN4 leucine zipper produced cells with a blue phenotype. The leucine zipper could be substituted both by the CC-Di homodimer and by the CC-Di-AB pairs to give the blue phenotype indicative of adenylate cyclase reconstitution. Moreover, the heterodimers produced a graded effect on phenotype: a strong blue phenotype was observed in strain DHM1.1 when both peptide sequences were at least 3.5 heptads long and reduced or no coloration seen when either partner was just 3 heptads long. The experiments were repeated in another *cya^-^* strain, BTH101, which is reported to be more sensitive to weak interactions (Figure 1c). In this strain a positive interaction in cells expressing CC-Di-B^3^ in conjunction with CC-Di-A^3.5^ or CC-Di-A^4^ was more evident, although the intensity of the blue phenotype was reduced in all cases compared to DHM1.1.

These results indicate that the homodimeric CC-Di and the heterodimeric CC-Di-AB pairs form PPIs within the cellular environment when expressed as fusions to a split cytoplasmic enzyme.

### Transcription activation *in vivo* by *de novo* designed PPIs

The graded phenotype of the heterodimeric adenylate cyclase constructs suggests that the binding affinities designed and measured *in vitro* are reflected in the strength of interaction *in vivo.* However, this adenylate cyclase assay is only semi-quantitative as it contains a positive feedback loop (expression of the fusion proteins is increased by the production of cAMP). To test the behavior of the CC-Di peptide sequences in a more quantitative system we next determined their ability to drive transcription activation in a bacterial 2-hybrid system.

Arbitrary PPIs can activate transcription via recruitment of RNA polymerase when one interacting partner is fused to a sequence-specific DNA binding domain (DBD) and the other is fused to RNA polymerase.^14^ To test the *de novo* CCs as transcription-activating interfaces we used a bacterial 2-hybrid system comprising the λcI repressor protein as the DBD and a truncated α subunit of RNA polymerase as the target for recruitment (Figure 2a).^30^ CC-Di, CC-Di-A or CC-Di-B sequences were fused to the *C* termini of the truncated α subunit and the DBD. The ability of combinations of these constructs to activate transcription was monitored in a reporter strain carrying a *lacZ* reporter gene under the control of a synthetic promoter with an upstream λcI binding site.

**Figure 2.**
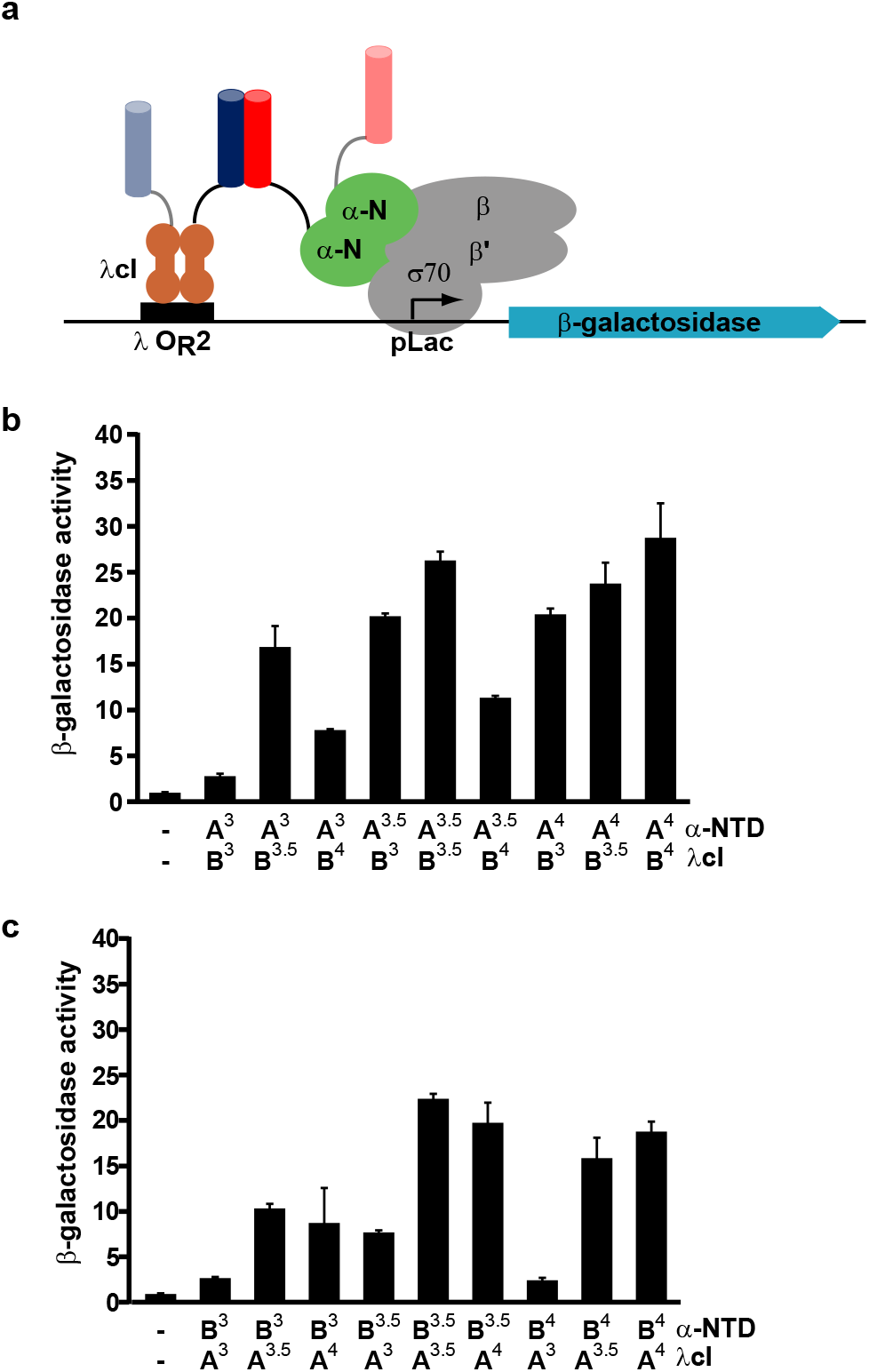
Activation of gene expression with *de novo* designed PPIs. (a) In this assay one CC peptide is fused to the λCI protein and the other is fused to the NTD of the α subunit of RNAP. Formation of a CC recruits RNAP to the *lac* promoter which activates transcription of β-galactosidase. (b & c) Bar charts of β-galactosidase activity of cells expressing fusion proteins containing different combinations of the 3-, 3.5- and 4-heptad repeat heterodimeric CC peptides. The acidic coils (CC-Di-A) were fused to the α-NTD and the basic coils (CC-Di-B) were fused to λcI, and *vice versa.* β-galactosidase activity was normalized to the OD_600_ of the bacterial cell culture and is the average of activity from three different cultures shown with standard error.

The homodimeric CC-Di fusions did not activate gene expression in this system (Supplementary Figure S1). This is not surprising as both λcI and the α subunits are themselves dimers, so we expect only *in cis CC* homodimerization and no *in trans* DBD-target interactions. By contrast, all the CC-Di-AB combinations activated transcription, regardless of which of the AB pairing was fused to the α subunit and which was fused to the DBD (Figure 2 b and c). Activation depended on the presence of a cognate binding partner (Supplementary Figure S1), and in any given orientation the degree of activation increased with the length of the PPI for combinations of sequences containing 3 and 3.5 heptad repeats. Activation by CC-Di-AB combinations in which one or both partners contained a 4-heptad repeat showed less predictable levels of transcription activation. At present we cannot offer a clear explanation of this, although it may reflect competition between on-target heterodimerization and off-target homodimerization of these longer CCs in a manner similar to that seen with CC-Di. This unexpected complexity highlights the need for some empiricism in the use of these *de novo* designed systems. Nonetheless, it is clear that gene activation can be directed by these *de novo* designed heterodimeric PPIs *in vivo.*

### Transcription repression *in vivo* by *de novo* designed PPIs

The *E. coli* Lac repressor (LacI) is a “dimer of dimers”, with the primary dimer interfaces between monomer surfaces and tetramerization mediated by C-terminal regions of each monomer, which form an antiparallel four-helix CC bundle.^31^ Dimerization enables the repressor to bind tightly to a palindromic operator sequence, and tetramerization enables simultaneous binding to a second, auxiliary operator.^32^ The CC region can be replaced by the GCN4 leucine zipper, converting the tetrameric protein into an active dimer.^31^ Similarly, we replaced the wild-type CC with our *de novo* designed CC dimers (Figure 3a). To maximize the reliance of dimerization on interaction of our CC sequences we used a C-terminally truncated LacI variant with a weakened monomer-monomer interface (LacI*).^33^

**Figure 3.**
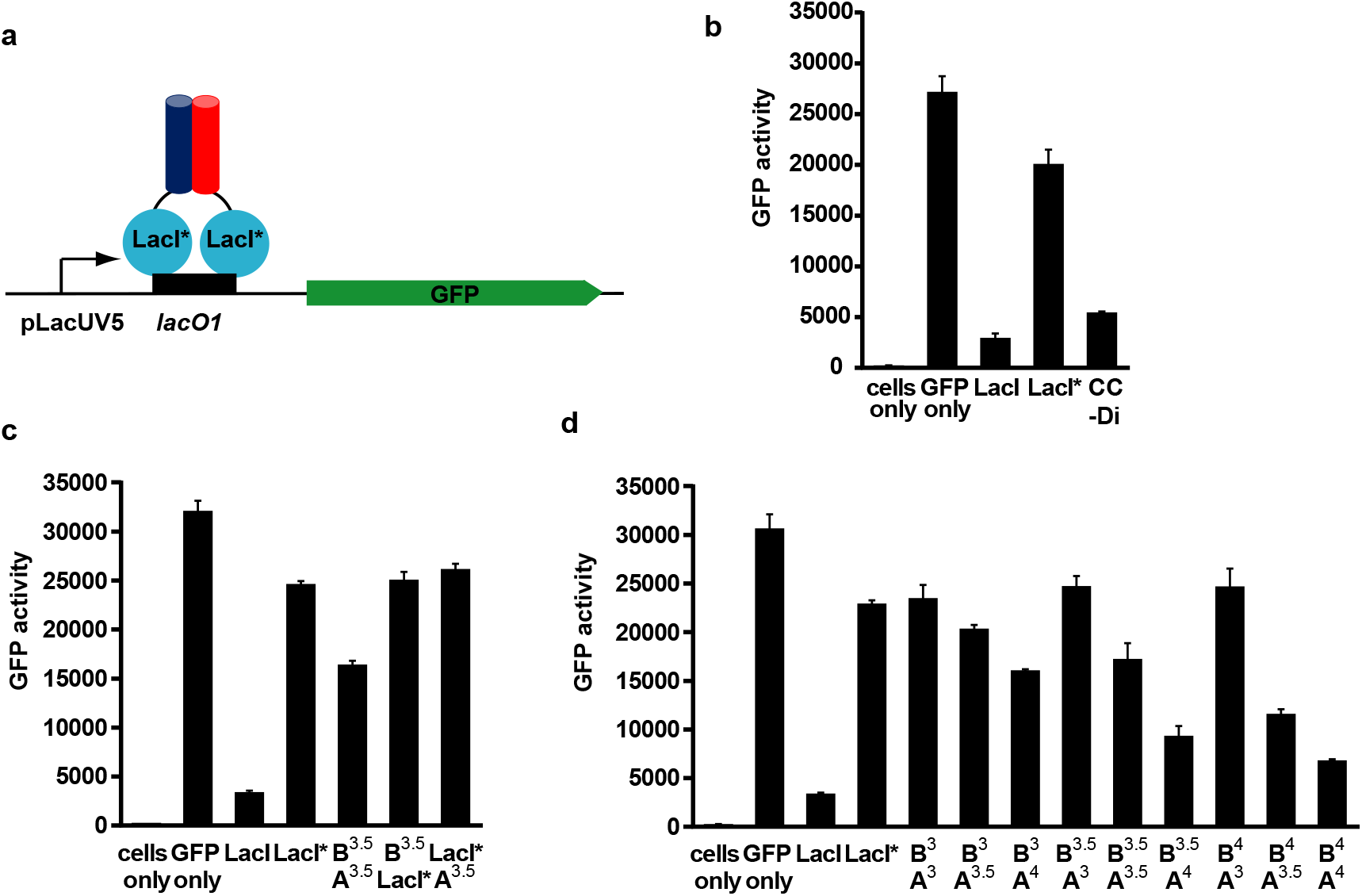
Repression of transcription mediated by interaction of *de novo* designed PPIs. (a) CC peptides were fused to LacI*, a dimerization mutant of Lac repressor. If interaction of the CC peptides occurred LacI* was able to bind to *lacO1* and repress transcription of GFP. (b) Bar chart showing repression of GFP activity mediated by the interaction of homodimeric coiled coil peptides (CC-Di) fused to LacI*. (c) Bar chart of GFP activity when LacI* was fused to either an acidic or basic 3.5 heptad repeat heterodimeric CC peptide (CC-Di-A^3.5^ or CC-Di-B^3.5^) and assayed in the combinations indicated. (d) Bar chart showing repression of GFP activity of cells expressing LacI* fusion proteins containing different lengths of the heterodimeric CC peptides CC-Di-A and CC-Di-B. GFP fluorescence was normalized to the 0D600 of the cell culture and is an average of three repeats shown with standard error.

The *de novo* CCs were fused *via* short linkers to the *C* terminus of LacI^*^. Genes encoding full length LacI or the LacI* proteins, with *N*-terminal His-tag and Xpress^TM^ tags, were expressed under the control of the arabinose-inducible P_BAD_ promoter. Activity of the resulting fusion proteins was tested in a *lacl*^-^ strain using a superfolder GFP reporter under the control of the *lacUV5* promoter, which carries a single *lac* operator sequence (Figure 3a). GFP expression was greatly reduced in cells expressing full length LacI protein, but only slightly reduced in cells expressing LacI* (Figure 3b). GFP expression in cells expressing LacI*-CC-Di was similar to that observed with full length LacI, indicating that CC-Di can substitute for the WT CC sequence to drive oligomerization of the repressor protein and consequent binding to DNA.

Next, we measured the effect of forming LacI* heterodimers mediated by the tunable CC-Di-AB series. The LacI*-CC-Di-A and LacI*-CC-Di-B constructs were expressed from different plasmids, each under the control of the P_BAD_ promoter. Co-expression of LacI* proteins fused to CC-Di-A^3.5^ and CC-Di-B^3.5^ resulted in a level of repression that was intermediate between full-length LacI and LacI* (Figure 3c). This effect requires a complementary partner sequence: neither LacI*-CC-Di-A^3.5^ or LacI*-CC-Di-B^3.5^ increased repression compared to LacI* when expressed without its partner. Notably, the series of CC-Di-AB fusion proteins repressed expression of GFP in line with the affinities of the CC heterodimers measured *in vitro* (Figure 3d).^24^ For example, cells expressing LacI*-CC-Di-B^3.5^ showed stronger repression when co-expressed with a fusion partner carrying a 3.5-heptad CC-Di-A sequence than they did when paired with a 3-heptad variant, and the level of repression increased further when the 4-heptad CC-Di-A sequence was used. The pattern of increased repression with increasing predicted strength of CC interaction was observed with all of the tested combinations, with the strongest repression being observed with the pairing of the two 4-heptad repeat sequences.

These results confirm that the homodimeric CC-Di and the heterodimeric CC-Di-AB modules can function as PPIs to mediate the affinity of dimerization of transcriptional repressors *in vivo* in a predictable and tunable fashion.

### Oligomerisation of TAL-based repressors by *de novo* designed PPIs

To create truly orthogonal synthetic transcription repressors it is desirable to couple designed PPIs with designable DBDs. CRISP-Cas9, Zn-fingers and TAL repeats have all been used to direct protein binding to specific sites on DNA within cells.^34,35^ TAL effector proteins (TALEs) contain tandem arrays of ≈34-residue TAL-repeat DBDs, each of which recognizes a single target base in DNA.^36^ Site-specific DNA binding proteins can thus be built by assembling appropriate combinations of these TAL repeats. As a step towards creating wholly designed systems in which the specificity and affinity of both protein-protein and protein-DNA interactions can be specified, we combined our *de novo* CC-based PPIs with engineered TAL-based DBDs to create tunable homodimeric and heterodimeric transcription factors.

In their natural context the arrays of TAL repeats are flanked by *N-* and C-terminal regions that appear to be important for function in mammalian cells.^37^ To identify the minimal TAL repeat scaffold that can serve as a DBD in our bacterial system, we designed a series of TAL-repeat proteins to bind to the *lacO1* operator sequence; namely, (I) a full-length TALE protein with intact *N-* and *C*-terminal regions, and truncated proteins lacking (II) the *N*-terminal region, (III) the C-terminal region, or (IV) both. These were expressed from the P_BAD_ promoter, and their ability to bind DNA *in vivo* was assessed with a GFP reporter gene expressed from the *lacUV5* promoter carrying a single copy of *lacO1* (Figure 4). Construct I repressed the reporter gene efficiently at basal and induced levels of expression. Construct IV produced no repression at any level of expression tested. Construct II showed substantially impaired repression, although some function was retained. In contrast, although construct III was less effective than the full-length protein it did repress transcription effectively when its expression was induced.

**Figure 4.**
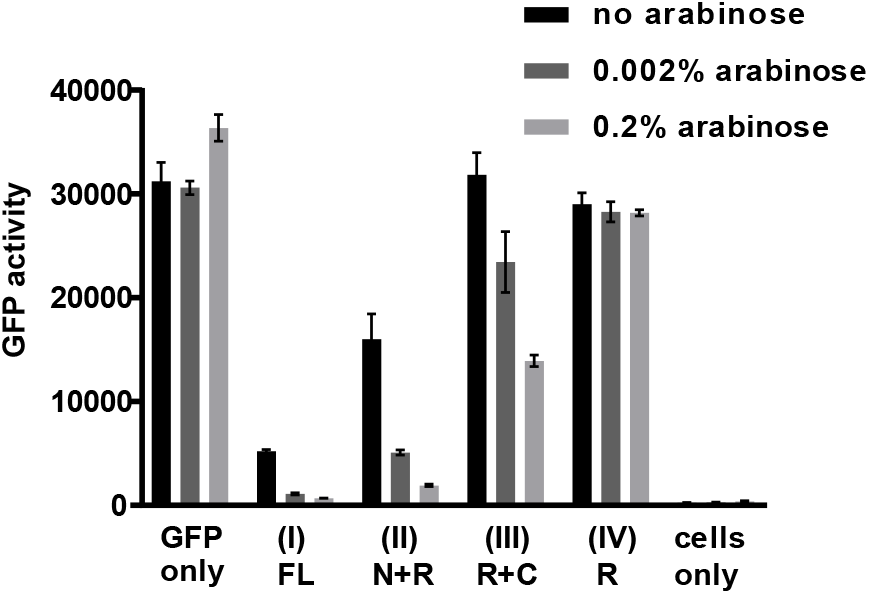
Repression of GFP activity by full-length and truncated TALE proteins. Cells were transformed with plasmids expressing either a full-length TALE (I), or derivatives lacking the *C*-terminal region (II), the *N*-terminal region (III) or both the *N-* and *C*-terminal region (IV). N: N-terminal region. R: TAL repeat region. C: C-terminal region. Arabinose was added to the cells at the concentrations indicated in order to induce expression of the TAL protein. GFP fluorescence was normalized to the OD_600_ of the cell culture and is an average of three repeats shown with standard error.

As the C-terminal region of the TALE protein is not essential for DNA binding function in bacteria, we fused the homodimeric CC-Di peptide sequence *via* a short linker to the *C* terminus of a 17-repeat TAL array that was designed to bind to the *lacO1* sequence and that retained the native N-terminal region (TALX). Dimeric TALE-based proteins can loop DNA, enhancing the efficiency of repression.^12^ Therefore, we tested the ability of this construct to repress transcription from *lacUV5* promoters carrying either one or two *lacO1* sequences. Each contained a “primary operator” that overlapped the transcription start site, and the second operator, when present, was placed 92 bp upstream of the primary operator (Figure 5a). Control experiments with wild-type tetrameric LacI confirmed that the presence of the auxiliary upstream *lacO1* promoter enhanced repression in our system (Figure 5b). TALX lacking CC-Di repressed transcription from the single and dual operator promoters equally, but repression by TALX-CC-Di was enhanced by the presence of the upstream operator. This enhancement was abolished when the sequence of the upstream operator was changed from that of *lacO1* to the related but distinct sequence, *lacO3.* These results suggest that TALX-CC-Di forms a dimer *in vivo* that, by looping DNA, can bind cooperatively to two specified DNA sites.

**Figure 5.**
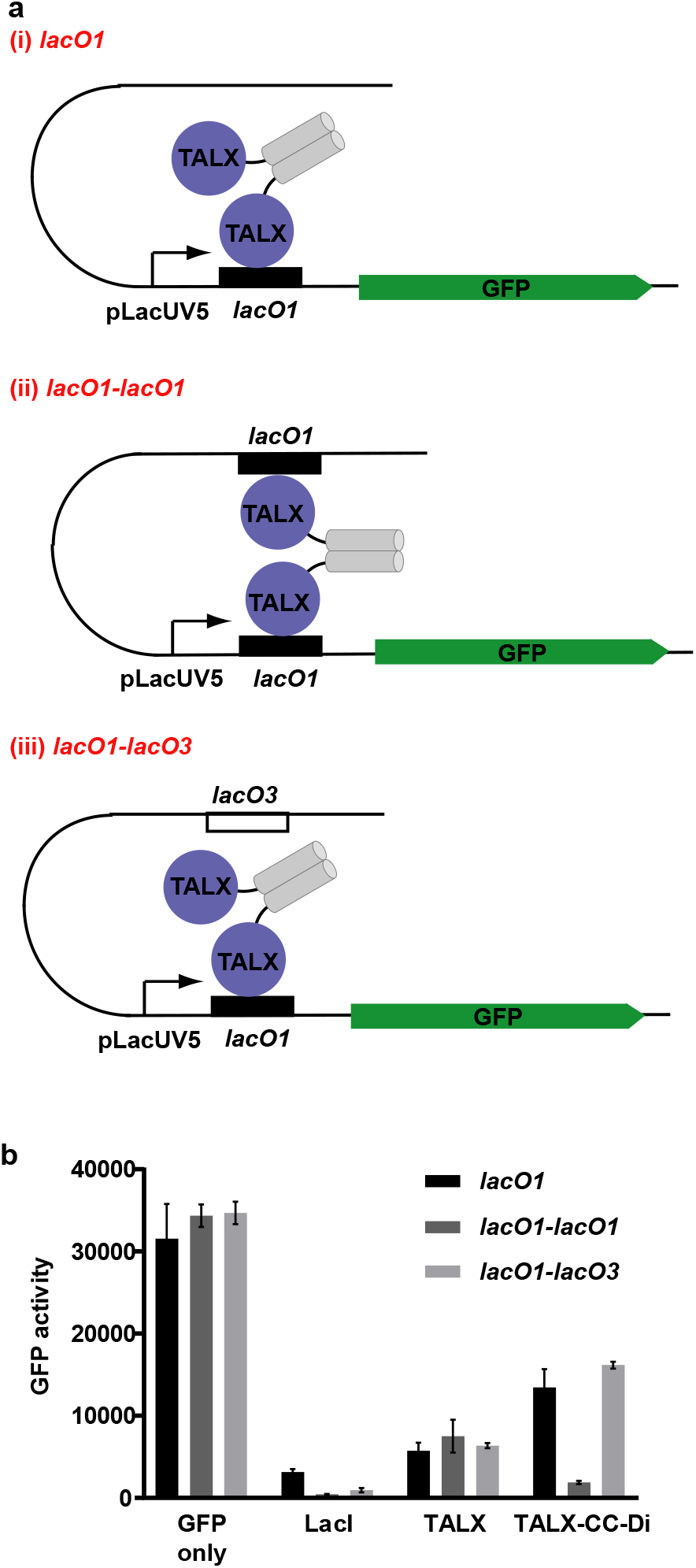
Repression of GFP activity by *de novo* homo-dimeric TAL-CC fusion proteins. (a) CC-Di was fused to TALX which binds the *lacO1* operator. Three GFP reporter plasmids were used in which there was (i) one *lacO1* site at the promoter, (ii) two *lacO1* sites 92 bp apart, and (iii) the upstream binding site was changed to the operator sequence *lacO3.* (b) Bar chart showing GFP activity of cells expressing the GFP reporter plasmid and the repressor construct indicated. Repression of GFP was enhanced when two binding sites for TALX were present and the repressor protein was able to dimerize via CC-Di. GFP fluorescence was normalized to the OD_600_ of the cell culture and is an average of three repeats shown with standard error.

Heterodimerization of TAL constructs should allow looping between two different DNA sequences, and also offer the possibility of integrating multiple regulatory signals to control the expression of each partner. To test this, we combined TAL constructs with the CC-Di-AB heterodimerization system. We designed a second 16 repeat TAL array, which retained the native *N*-terminal region and bound a target site that was not recognized by TALX (TALY) (Supplementary Figure 2). We fused CC-Di-B^4^ *via* a short linker to the *C* terminus of TALX and CC-Di-A^4^ *via* a short linker to the *C* terminus of TALY. Then, we tested the effect of combinations of constructs on expression from *lacUV5* promoters carrying the TALY binding site as a primary operator and *lacO1* or *lacO3* as the secondary operator (Figure 6). Coexpression of TALY-CC-Di-A^4^ and TALX-CC-Di-B^4^ enhanced repression when the auxiliary operator was *lacO1.* This enhancement was lost when the upstream site was mutated to *lacO3,* or when the PPI was abolished by omission of the CC-Di-A/B peptide from TALX or TALY.

**Figure 6.**
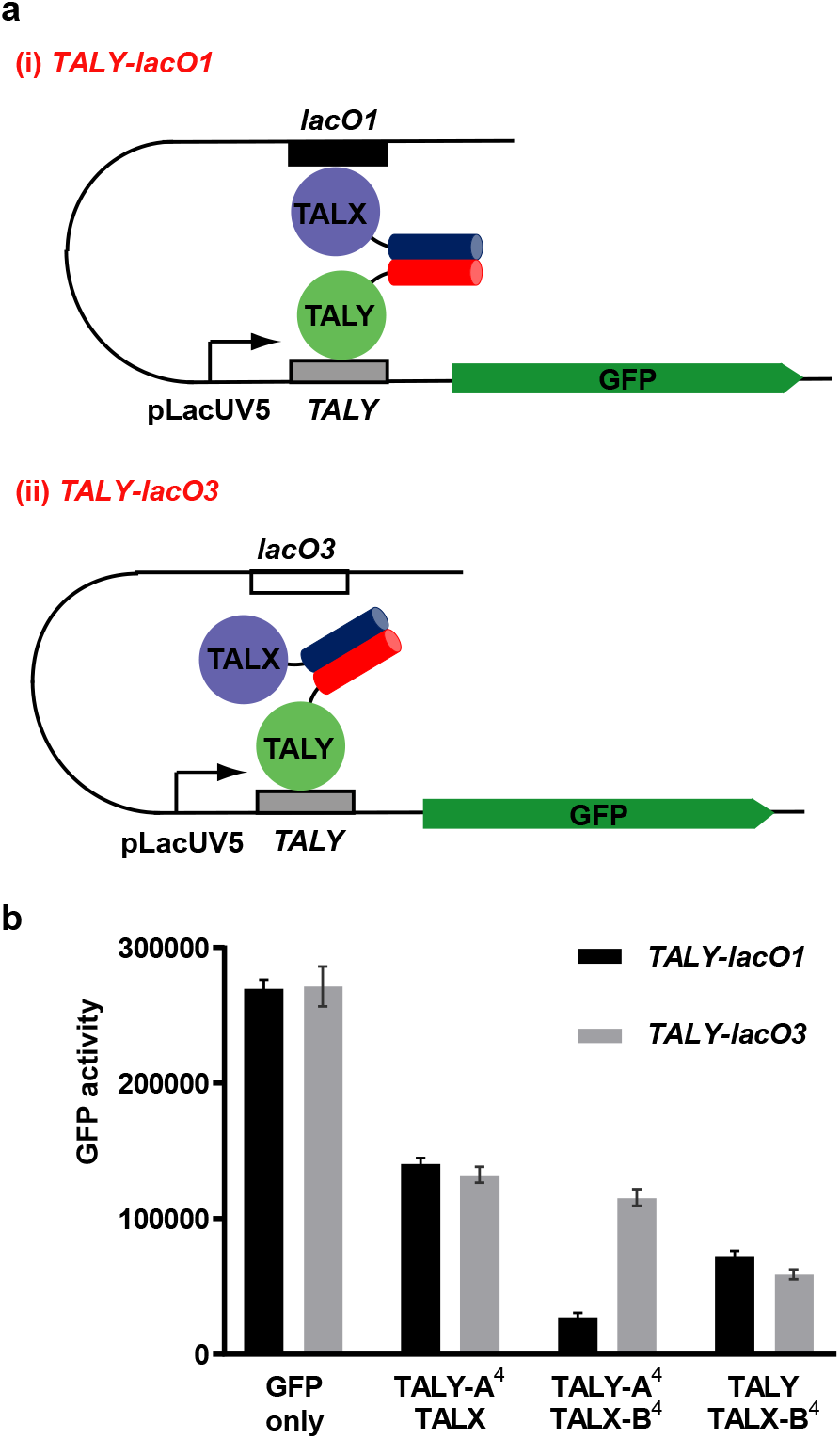
Repression of GFP activity by *de novo* heterodimeric TAL-CC fusion proteins. (a) CC-Di-A^4^ was fused to TALY and CC-Di-B^4^ was fused to TALX. (i) A GFP reporter plasmid was used in which there was a TALY binding site at the promoter and a *lacO1* site 92 bp upstream. (ii) An additional reporter plasmid was used where the upstream binding site was changed to the *lacO3* sequence. (b) Bar chart of GFP activity of cells transformed with a GFP reporter plasmid and two additional plasmids expressing the TALX and TALY fusion proteins as indicated. Repression of GFP was enhanced when binding sites for TALX and TALY were present and the repressor protein was able to dimerize via CC-Di-A^4^B^4^ interactions. GFP fluorescence was normalized to the OD_600_ of the cell culture and is an average of three repeats shown with standard error.

These results indicate that combining TAL repeat sequences with *de novo* designed PPIs allows the design of proteins with desired protein-DNA and protein-protein interaction specificity that function within living cells.

### Conclusion

The ability to direct and control the assembly of macromolecular complexes in cells is a key aim of synthetic biology. For instance, building networks of interacting components could allow engineered cells to colocalize or to segregate cellular processes, and to respond to their environment in complex but predictable ways. Herein, we show that straightforward *de novo* designed protein-protein interactions (PPIs) can substitute for natural PPIs to complement fragments of enzymes and to control transcriptional processes in bacterial cells. In addition, by combining these *de novo* PPIs with engineered DNA-binding repeats, we generate completely new transcriptional repressors. Moreover, because of the designability of the *de novo* PPIs, the degree of downstream activity can be tuned. These *de novo* and engineered modules expand the repertoire of components for synthetic biology and protein design in the cell.

The construction of Gene Regulatory Networks (GRNs) is one area where multiple orthogonal and tuneable PPIs of the type we describe are needed. In this field, transcription repressors and activators, together with their DNA targets, are organized in topologies that enable cells to undertake computational tasks and actuate appropriate responses.^38,39^ Some of the most complex GRNs have been built in *E. coli,* where a wide range of well-characterized native components is available. However, as the complexity of the networks increases the use of endogenous regulatory components becomes limiting. Many of the existing GRN subsystems reuse the same small set of transcription factors, such as the LacI and TetR repressors.^40,41^ Therefore, they cannot be combined readily as cross-talk between different parts of the network is inevitable. The range of characterized components available for use in GRNs can be increased either by co-opting regulatory components from other organisms, or by creating novel components. An example of the first approach is a library of mutually orthogonal repressors composed of TetR proteins from diverse prokaryotic species.^42^ New components can also be created by modifying existing natural systems to modify their properties and make them orthogonal; for example, mutation of the bacteriophage T7 RNA polymerase has been used to generate a library of orthogonal RNA polymerases that recognise different promoter sequences.^43^ In addition, transcription regulators represent an attractive target for *de novo* protein design, which was part of the motivation for the work presented herein.

Here we show that *de novo* CC-based PPIs designed from first principles can mediate the function of both transcription activators and repressors. Furthermore, these *de novo* PPIs can be combined with engineered TAL DNA-binding repeats to produce transcription repressors in which the affinity and specificity of both protein-protein and protein-DNA interactions are specified. This offers possibilities for creating components with specificities and affinities that are optimized on the basis of the mathematical model of a desired GRN, avoiding the limitations of natural components that have evolved for other purposes.

We have explored the function of a toolkit of designed homo- and heterodimeric CCs in four different molecular contexts in *E. coli* cells. We find that in most cases the CC behavior mirrors that seen *in vitro.* Some of the peptide sequences tested here have been shown recently to assemble in *E. coli* in other contexts: the heterodimeric CC drives the assembly of a novel cytoscaffold and the subcellular localization of active enzymes when fused to shell proteins of a bacterial microcompartment;^6,7^ and, while the work presented here was in preparation, the same heterodimeric CCs have been shown by others to recruit T7 RNA polymerase to Zn-finger DNA-binding domains.^28^ Some adverse context-dependent effects have been noted: in our activation experiments proximity effects may inhibit heterodimerization; and in the programmable T7 RNA polymerase system the hierarchy of CC interaction strength varies with the nature of the Zn-finger domains to which the peptides are fused.^28^ Thus, it is likely that improved rules or methods for designing linker sequences will be needed to help minimize such effects in future applications. We are working on this challenge using the ISAMBARD suite for computational protein design.^44,45^ Nonetheless, our results, together with those of others,^26,28^ indicate that the rules used to design our peptide sequences are sufficiently comprehensive to allow the CC components of the sequences to interact as designed in a cellular environment. Although we have yet to probe these systems with proteomics, it appears that the introduced biomolecular interactions operate orthogonally to the endogenous *E. coli* proteome and interactome. This work provides a starting point for the design and implementation of more-complex higher-order PPIs and possibly regulatable PPIs for control of protein assembly within cells.

## MATERIALS AND METHODS

### Plasmids

Full details of the construction of the plasmids used in this work are given in the supplementary information. Briefly: Adenylate cyclase reconstitution assays used derivatives of plasmid pKT25 *(kan^R^),* which encode fusions to the T25 fragment of *Bordetella pertussis* adenylate cyclase (CyaA) (amino acids 1-224) and of plasmid pUT18c *(amp^R^),* which encode fusions to the T18 fragment of CyaA (amino acids 225-399).^46,47^ Transcription activation assays used derivatives of pRA02 *(amp^R^),* which encodes fusions with the α-subunit of RNA polymerase (amino acids 1-248), and of pRA03 *(cm^R^)* which encodes fusions with the λcI protein (amino acids 1-236)^48^. Lac repressor protein fusions were expressed from derivatives of plasmid pBADLacI* (*amp*^R^) or pVRcLacI* (*cm*^R^) which encode a *C*-terminally truncated Lac repressor (amino acids 1-331) containing an L251A substitution, under the control of the arabinose inducible *araBAD* promoter. Fusions to TAL repeats were expressed from plasmids derived from pVRc20_992 *(cm^R^,* a gift from Christopher Voigt, Addgene #49739^49^) or pBADHis-B-iRFP (a gift from Vladislav Verkhusha, Addgene plasmid #3 1 8 5 5^50^), under the control of the arabinose inducible *araBAD* promoter. The reporter plasmid pVRbLacUV5 *(kan^R^)* and derivatives allow the expression of sfGFP from the *lacUV5* promoter and is derived from the plasmid pVRb20_992 *(kan^R^*, a gift from Christopher Voigt, Addgene plasmid #49714^49^).

### Bacterial two-hybrid assay utilizing adenylate cyclase reconstitution

The bacterial two-hybrid assay based on adenylate cyclase reconstitution described in this work is essentially that described by Battesti and Bouveret.^46^ cya-DHM1.1 or BTH101 cells^46^ were transformed with both pUT18c and pKT25 derived plasmids containing the adenylate cyclase subdomains T18 or T25 fused to different CC peptides. Cells were grown at 30°C on LB agar supplemented with 100 μg/ml ampicillin and 50 μg/ml kanamycin. Overnight cultures were diluted in LB to an OD_600_=1 and 2 μl of each culture was spotted onto LB agar + 100 μg/ml ampicillin + 50 μg/ml kanamycin + 0.5 mM IPTG + 40 μg/ml X-gal. Plates were incubated at 30°C for 24 hours (BTH101) or 48 hours (DHM1.1).

### Bacterial two-hybrid assay utilizing transcription activation

The transcription activation based bacterial two-hybrid assay described here is essentially that developed by Dove and Hochschild.^30^ Reporter strain KS1^14^ contains a *lacZ* gene on the chromosome with a promoter that can be activated by interactions between a peptide fused to λcI and a peptide fused to the α-subunit of RNA polymerase. KS1 cells were transformed with pRA02 and pRA03 or their derivatives and grown at 37°C on LB agar supplemented with 100 μg/ml ampicillin, 50 μg/ml kanamycin and 25 μg/ml chloramphenicol. Colonies were picked in triplicate and overnight cultures were grown at 37°C. These were used to inoculate 10 ml LB + 100 μg/ml ampicillin + 50 μg/ml kanamycin + 25 μg/ml chloramphenicol + 20 μM IPTG. Cultures were grown at 37°C until they reached an OD_600_ ~0.5. β-galactosidase activity of each culture was assayed in duplicate in 96-well plates after lysis by PopCulture lysis reagent (Novagen) essentially as described by Thibodeau *et al*.^51^ The change in A405 at 30°C was measured over 30 minutes at 1 minute intervals in a Spectramax plate reader (Molecular Devices) and the rate of change of the A405 was normalised by dividing by the OD_600_ of the cell culture.

### GFP assays

To monitor repression of transcription TB28 cells (MG1655ΔLacIZYA^52^) were transformed with pVRbLacUV5 reporter plasmid or its derivatives, and plasmids expressing Lac repressor or TALE fusion proteins as indicated. Colonies were picked in at least triplicate and overnight cultures were grown at 37°C in M9 minimal media + 0.25% glycerol + 0.5 mM CaCl2 + 2 mM MgSO4 + 2 μg/ml thiamine + 0.2% casamino acids (+ 50 μg/ml kanamycin + 100 μg/ml ampicillin + 25 μg/ml chloramphenicol where required). The overnight cultures were used to inoculate 10 ml of the same medium and cultures were grown at 37°C until they reached an 0D600~0.5. Where indicated arabinose was added to the 10 ml cultures at the concentrations indicated: where no arabinose was added the fusion protein expression resulted from basal transcription from the *araBAD* promoter. 5 ml of culture was centrifuged and the pellet was resuspended in 250 μl PBS (137 mM NaCl, 2.7 mM KCl, 10 mM Na2HP04, 2 mM KH2PO4). 2 x 100 μl cell suspension from each culture was placed in a black, flat bottomed 96 well plate and the fluorescence read in a FLEXstation plate reader (Molecular Devices). The excitation wavelength was 470 nm and the emission wavelength was 510 nm with a cutoff of 495 nm. GFP fluorescence (relative fluorescence units) was normalised by dividing by the 0D600.

## ACKNOWLEDGEMENTS

We thank Luke Cartlidge and George Black for technical assistance with the creation of TAL constructs and members of the DNA-protein Interactions Unit at the University of Bristol for helpful discussions. The authors are grateful to the BBSRC and EPSRC for funding through the BrisSynBio Synthetic Biology Research Centre (BB/L01386X1). D.N.W. holds a Royal Society Wolfson Research Merit Award (WM140008).

## Plasmid construction

The CC sequences and corresponding DNA sequences used in this work are shown in table S1. A short linker was encoded upstream of each CC peptide (table S2), and XbaI and Acc65I restrictions sites at each end facilitated cloning in frame with the C-terminal end of the various proteins used in this work. The DNA fragments encoding CC-Di-A^3.5^, CC-Di-B^3.5^ and CC-Di were synthesised as GeneStrings (GeneArt, Invitrogen). DNA sequence encoding CC-Di-A^3^, CC-Di-B^3^, CC-Di-A^4^ and CC-Di-B^4^ were produced by removing (CC-Di-A^3^, CC-Di-B^3^) or adding (CC-Di-A^4^, CC-Di-B^4^) sequences from/to expression vectors containing the 3.5 heptad CC sequences using PCR.

Plasmid pKT25 *(kan^R^)* encodes the T25 fragment of *Bordetella pertussis* adenylate cyclase (CyaA) (amino acids 1-224) and plasmid pUT18c *(amp^R^)* encodes the T18 fragment of CyaA (amino acids 225-399).^1,2^ Both fragments are expressed from the *PLac* promoter. The synthetic DNA fragments encoding the CC peptides were cloned in frame downstream of the T25 or T18 subdomains at XbaI/Acc65I sites to make pKT25-CC-Di, pUT18c-CC-Di, pKT25-B^3.5^ and pUT18c-A^3.5^. Plasmids pKT25-zip and pUT18c-zip encode the T25 and T18 fragments fused to the yeast GCN4 leucine zipper.^2^

For the transcription-activation based bacterial two-hybrid assay the DNA fragments encoding the CC peptides were cloned into either pRA02 *(amp^R^),* which encodes fusions with the α-subunit of RNA polymerase (amino acids 1-248), or pRA03 *(cm^R^)* which encodes fusions with the λcI protein (amino acids 1-236)^3^. DNA fragments encoding the CC peptides CC-Di-A^3.5^, CC-Di-B^3.5^ and CC-Di were inserted into the XbaI/Acc65I sites of both pRA02 and pRA03 allowing in-frame fusions with the α-subunit or λcI.

Plasmid pBADLacI (amp^R^) was made as follows: DNA encoding WT Lac repressor protein (LacI) was amplified by PCR from pET21a (Novagen) and was cloned into pBADHis-B-iRFP (a gift from Vladislav Verkhusha, Addgene plasmid #31855^4^) at BglΠ/HindΠI sites. pBADLacI allows expression of LacI from the arabinose inducible P_BAD_ promoter, giving an N-terminal 6xHis tag and Xpress^TM^ epitope tag. Plasmid pBADLacI* encodes a truncated LacI gene (aa 1-331) containing an L251A substitution which was introduced by site-directed mutagenesis. XbaI and Acc65I sites were introduced downstream of the truncated LacI gene, allowing DNA encoding CC-Di and CC-Di-B peptides to be cloned in frame at the *C* terminus of LacI*. In order to express heterodimeric LacI-CC peptide fusion proteins an additional set of *cm^R^* plasmids were made containing a different origin of replication. The *lacI\** gene, P_BAD_ promoter and *araC* gene were excised from pBADLacI* at BsaI/NsiI restriction sites and cloned into pVRc20_992 (a gift from Christopher Voigt, Addgene #49739^5^) to produce pVRcLacI*. DNA fragments encoding CC-Di-A peptides were cloned into pVRcLacI* at XbaI/Acc65I sites. The p15A *ori* in pVRcLacI* has a lower copy number than the pBR322 *ori* in pBADLacI* which lacks the *rop* gene^6^ so the expression levels of LacI*-CC-Di-A and LacI*-CC-Di-B are expected to vary slightly.

TALE DNA binding domain arrays were constructed using the Joung lab REAL assembly TALEN kit, a gift from Keith Joung (Addgene kit # 1000000017^7^). This kit allows the production of DNA fragments encoding TALE repeat arrays using sequential restriction enzyme digestion and ligation. The DNA binding sites of the TALEs used in this work are shown in table S3. pBAD-His-JDS78 *(amp^R^)* contains the N- and C-terminal domains of the TALE and also the 0.5 TAL repeat which is at the C-terminal end of the TALE repeat array, and was made by PCR amplification of a DNA fragment encoding the TALE *N* and *C* terminus and the T 0.5 repeat from the plasmid JDS78 (from Addgene kit # 1000000017). This fragment was cloned into the BglII /HindIII sites of a pBAD-His-iRFP derivative in which the BsmBI site was mutated, creating pBAD-His-JDS78. Sequence encoding a TAL repeat array recognising 16 bp of the *lacO1* sequence constructed by REAL assembly (TALA) was inserted into pBAD-His-JDS78 at the BsmBI site to create pBADTALA (Construct I; TALA aa 1-763). In order to examine the minimal TAL domains required for DNA binding the TALA repeat array (R) was also inserted into the vectors pBAD-His-JDS78ΔNTD (Construct II; TALA aa 129-763), pBAD-His-JDS78ΔCTD (Construct III; TALA aa 1-707) and pBAD-His-JDS78ΔNTD+CTD (Construct IV; TALA aa 129-707). These contained different combinations of the TALE N- and C-terminal regions and were created by PCR amplification from JDS78 and insertion into pBAD-His-RFP.

Plasmid pBAD-His-JDS78XA contains the N-terminal domain of the TALE and the T 0.5 repeat (but not the C-terminal domain) and was constructed by PCR using primers that added an XbaI site and Acc65I site downstream of the 0.5 TAL repeat. A sequence encoding a TAL repeat array recognising 17 bp of the *lacO1* sequence (TALX) was inserted into pBAD-His-JDS78XA to create pBADTALX, and then sequences encoding the CC peptides CC-Di and CC-Di-B were cloned in frame with at the *C*-terminal end to produce pBADTALX-CC-Di and pBADTALX-CC-Di-B. To express heterodimeric TAL-CC peptide fusion proteins an additional set of plasmids with an alternative *ori* and marker were constructed. A BsaI/NsiI fragment from pBAD-His-JDS78XA containing the araC gene and the expression cassette encoding the N-terminal domain of the TALE and the T 0.5 repeat was inserted into pVRc20_992 to produce pVRcJDS78XA. A sequence encoding a TAL repeat array recognising 16 bp of *non-lacO* sequence (TALY) was inserted into pVRcJDS78XA as above to create pVRcTALY. DNA fragments encoding CC-Di-A peptides were inserted at XbaI/Acc65I sites to produce pVRcTALY-CC-Di-A.

The reporter plasmid pVRbLacUV5 *(kan^R^)* allows the expression of sfGFP from the *lacUV5* promoter and is derived from the plasmid pVRb20_992 (a gift from Christopher Voigt, Addgene plasmid # 49714^5^). DNA containing the *lacUV5* promoter minus the CRP half site (-53/+40) was amplified from the plasmid pSRLacUV5^8^ by PCR and was cloned into pVRb20_992 at BspHI and BamHI sites. For experiments analysing repression by TAL-CC fusion proteins the following reporter plasmids were created by modifying pVRbLacUV5: *pVRblacO1-lacO1, pVRblacO1-lacO3,* pVRbTALY-lacO1, pVRbTALY-lacO3. Details of the promoter region of these reporter constructs is shown in table S4. Synthetic DNA fragments carrying the promoters containing binding sites for TALX (*lacO1*) and TALY, and the *lacO3* operator sequence were inserted into pVRbLacUV5 at EcoRI/HindIII sites. The spacing between the operator sequences is identical to the wild type *lac* promoter (92 bp).

## Western blotting

To detect expression of coiled coil peptide fusion proteins, bacterial cultures were lysed in SDS-loading buffer (100 mM Tris-Cl pH 6.8, 4% (w/v) SDS, 0.2 % (w/v) bromophenol blue 20% (v/v) glycerol, 200 mM DTT) and run on an SDS–polyacrylamide gel of an appropriate percentage. Protein was transferred onto an immobilonP PVDF membrane (Millipore). Membranes were probed with polyclonal anti-α subunit antibodies (a gift from A. Ishihama), anti-λCI antibodies (a gift from A. Hochschild) or a monoclonal antibody against the His-tag (BD-biosciences #631212) using standard western blotting techniques. Detection was carried out using the POD chemiluminescence system (Roche).

**Table S1:**
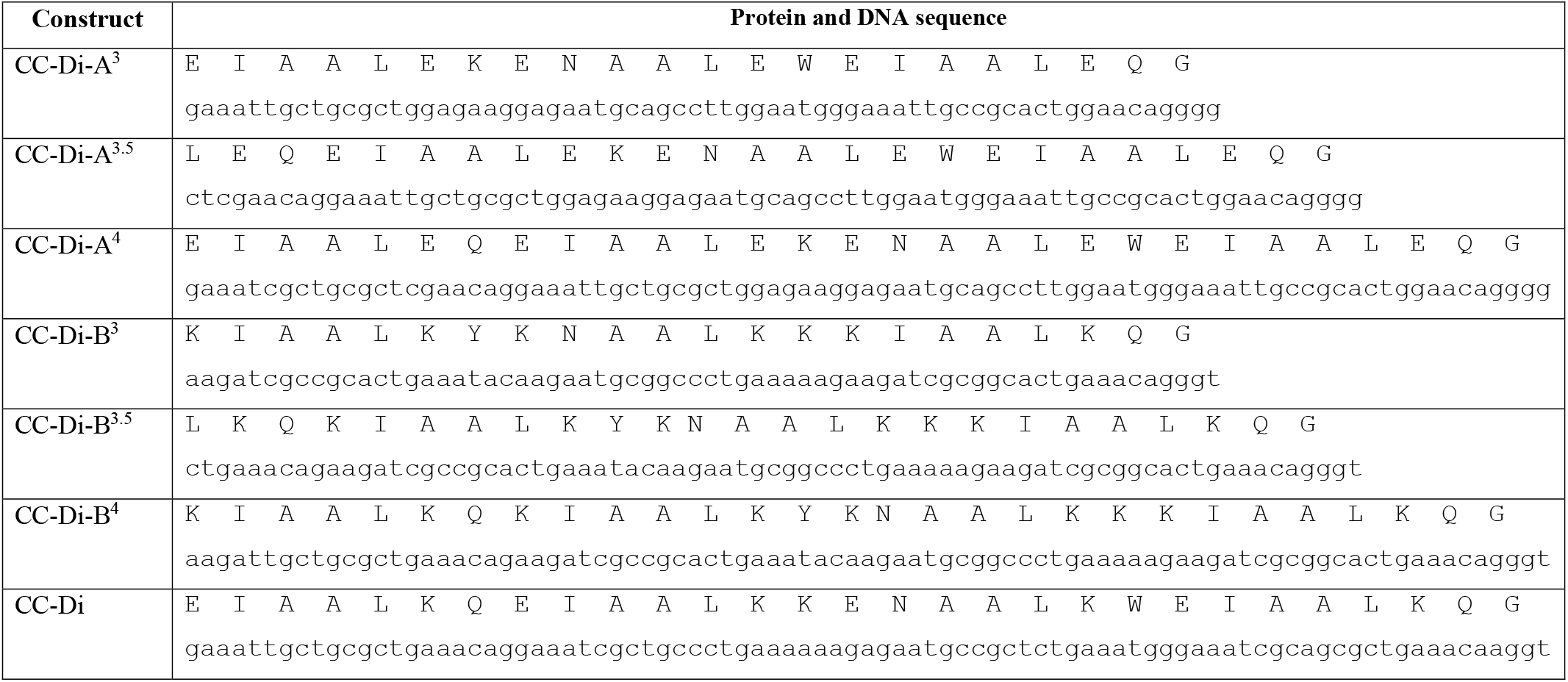
The amino acid sequences of the CC peptides used in this work^9,10^. Below is the DNA sequence which was designed using codons selected for expression in *E. coli.*

**Table S2:**
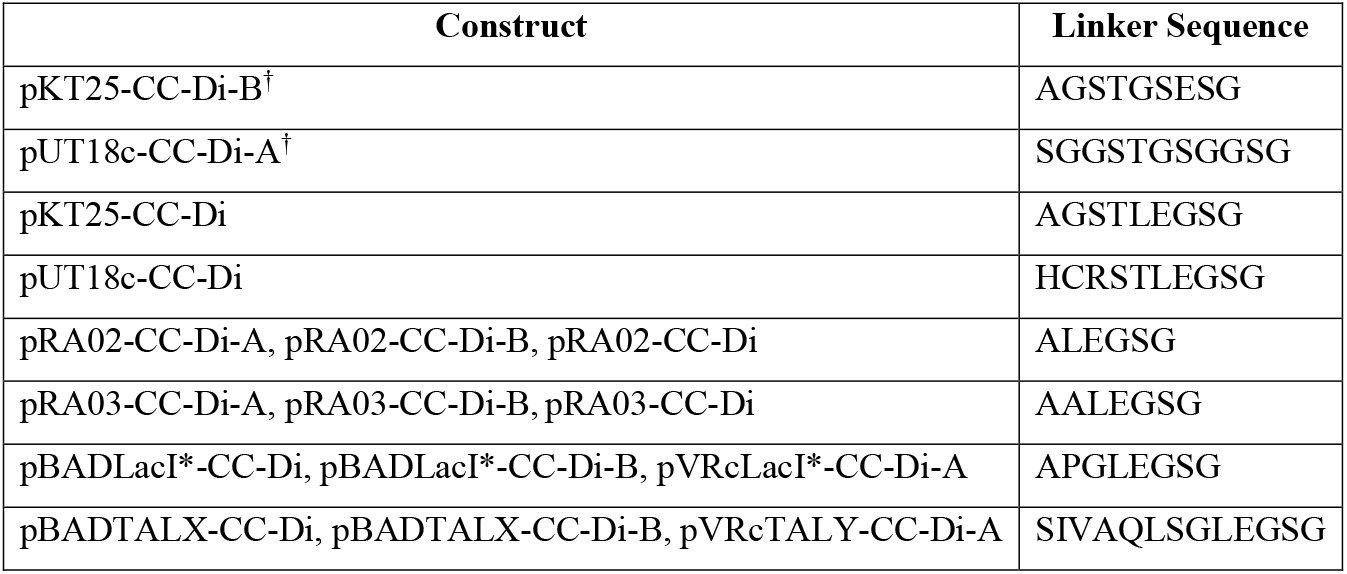
Plasmids used in this study and the linker sequences between the protein domains and the C-terminal CC peptides. ^†^These linker sequences were optimised by molecular dynamics modelling to allow optimal positioning of the CC and modified from the original sequence using site-directed mutagenesis.

**Table S3:**
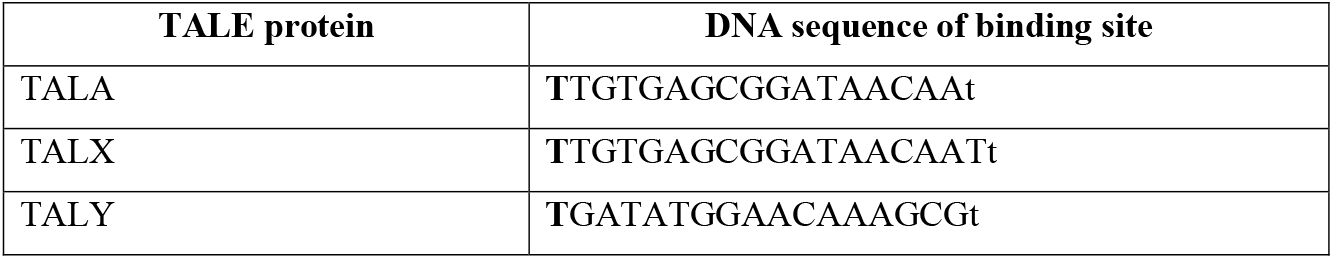
DNA Binding sites for the TALE proteins used in this work. The first T in bold is not bound by the TAL repeat array but is a requirement for TALE binding. The lowercase t is recognised by the 0.5 TAL repeat present at the C-terminal end of each array.

**Table S4:**
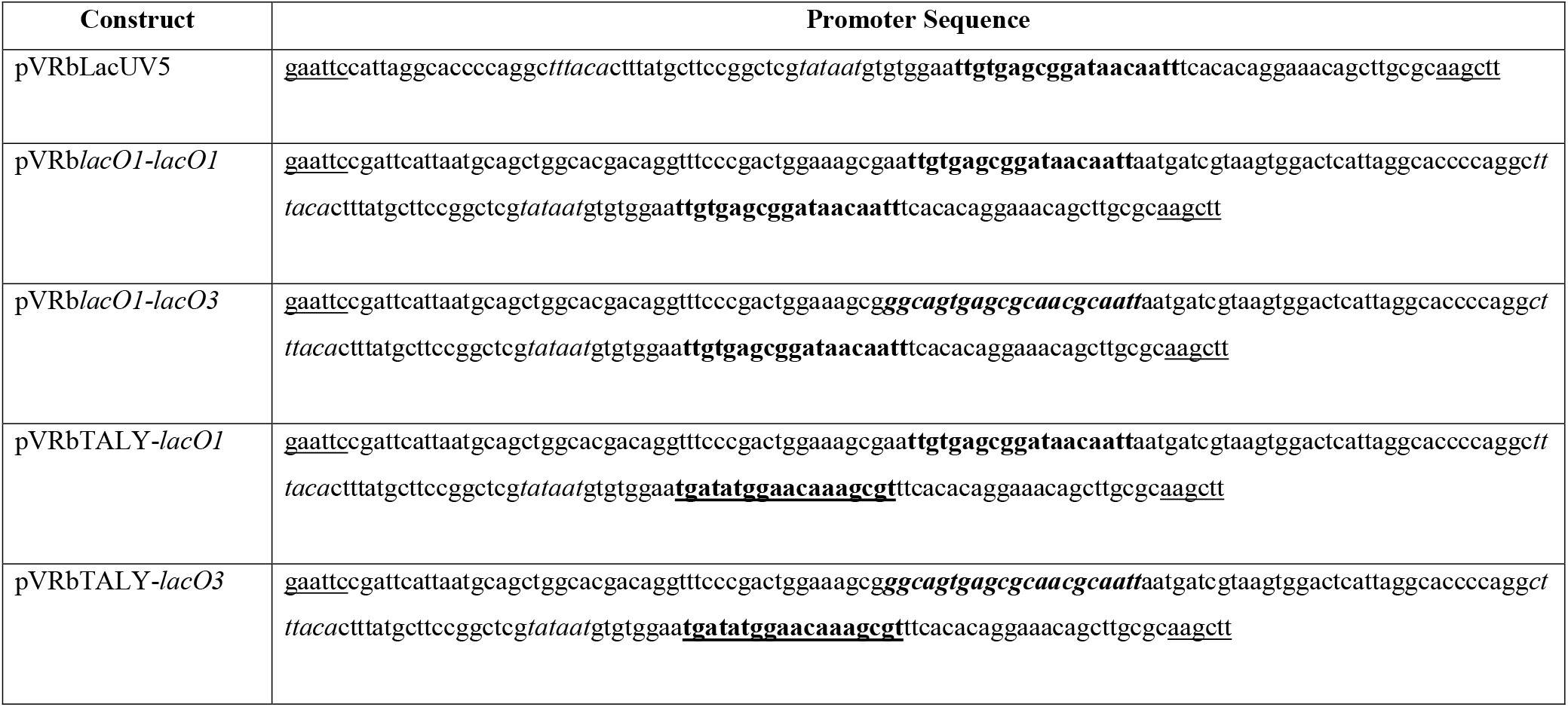
DNA sequences of the promoter regions of sfGFP reporter constructs. EcoRI and HindIII restriction sites are underlined. -35 and -10 sequences of the LacUV5 promoter are in italics. The TALX binding site *(lacO1)* is in bold, TALY is in bold and underlined and *lacO3* is in bold and italics.

**Supplementary figure S1.**
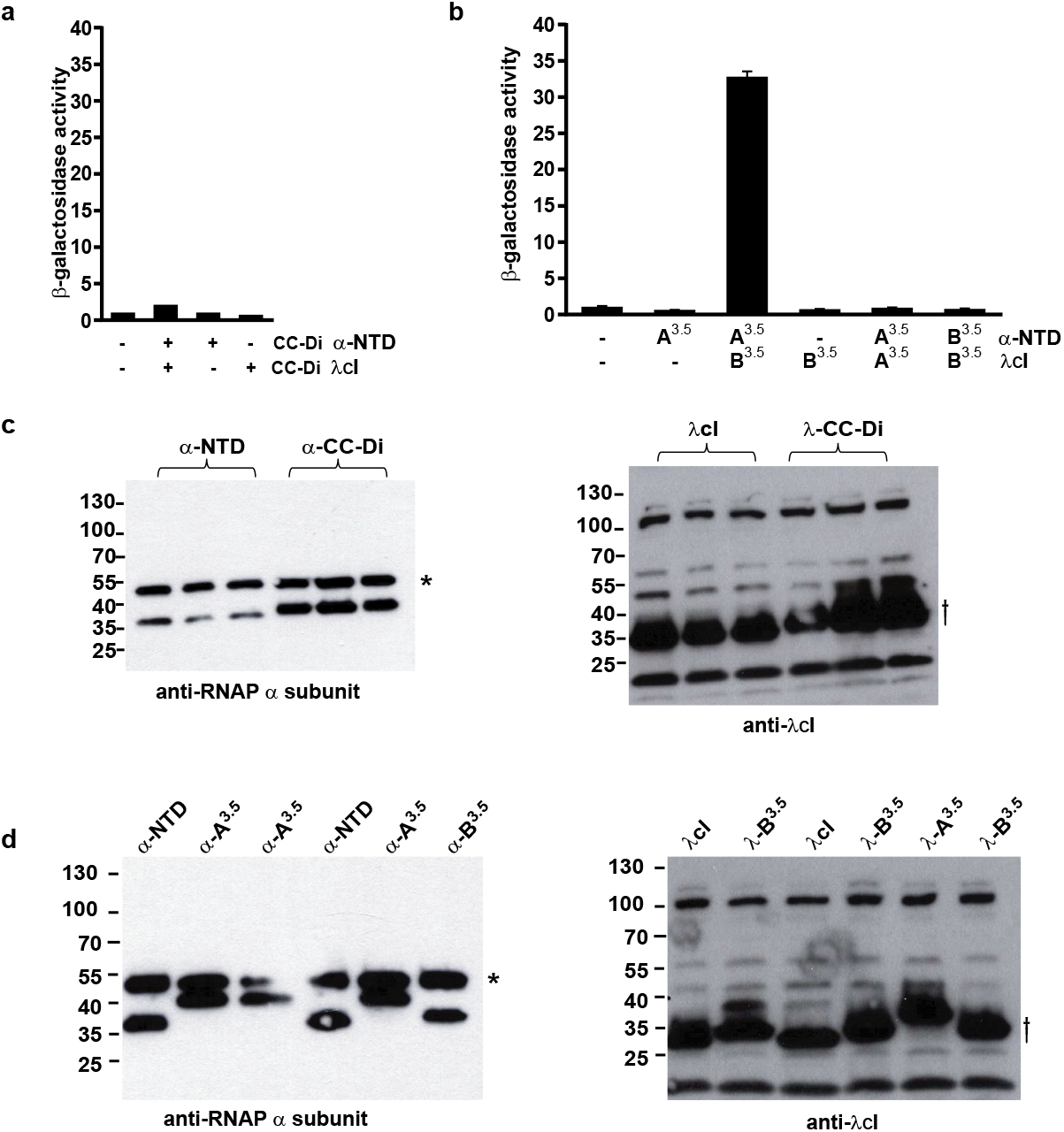
(a) Bar chart showing β-galactosidase activity of cells expressing fusion proteins containing CC-Di. Transcription activation is not observed with the homodimeric CC-Di. (b) Bar chart showing β-galactosidase activity of cells expressing fusion proteins containing CC-Di-A^3.5^ and CC-Di-B^3.5^. Transcription activation only occurs when both peptides of the CC are present. β-galactosidase activity was normalised to the OD_600_ of the bacterial cell culture and is the average of activity from three different cultures shown with standard error of the mean. (c & d) Western blots showing expression of fusion proteins in the assays shown in a & b. Blots were probed with an antibody against the α-subunit of RNAP (* indicates the cellular α-subunit which is also recognised by the antibody) and an antibody against the λCI protein (f shows the λCI fusion protein, the other bands are likely to be nonspecific products). (c) Blot showing expression of fusion proteins from cultures assayed in (a). Lanes 1-3 are three separate cultures expressing α-NTD of RNAP and λCI protein alone. Lanes 4-6 are three separate cultures expressing α-NTD-CC-Di and λcI-CC-Di fusions. (d) Blot showing expression of fusion proteins from cultures assayed in (b). Each lane of the gel is cell lysate from one of the three cultures assayed. CC-A^3.5^ and CC-B^3.5^ were fused to the α-NTD and to λcI in different combinations as indicated.

**Supplementary figure S2.**
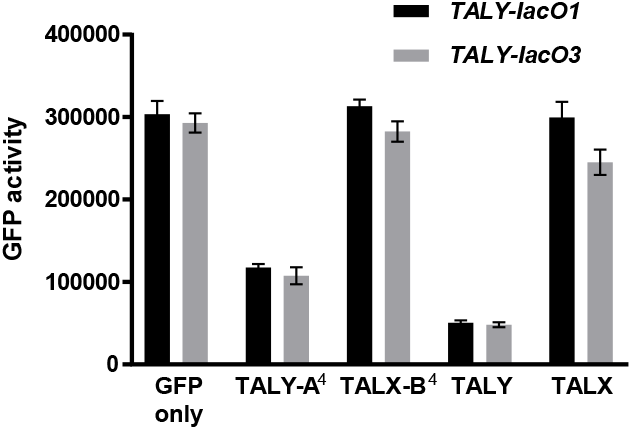
Control for the experiments in figure 6, repression of GFP activity by *de novo* heterodimeric TAL-CC fusion proteins. Bar chart showing GFP activity of cells transformed with GFP reporter plasmids and with a plasmid expressing single TALX or TALY fusion proteins as indicated. Where shown CC-Di-A^4^ was fused to TALY protein and CC-Di-B^4^ was fused to TALX protein. GFP fluorescence was normalised to the OD_600_ of the cell culture and is an average of three repeats shown with standard error.

AUTHOR CONTRIBUTIONS
AJS, DNW and NJS designed the study, interpreted results and wrote the paper. AJS conducted all experimental work. FT designed sequences for the expression of CC peptides. DS designed linkers for fusion of CC peptides to other proteins. All authors commented on the final draft of the paper.

## REFERENCES

1. Boyle, A. L.; Woolfson, D. N., De novo designed peptides for biological applications. Chem Soc Rev 2011, 40 (8), 4295–306.

2. Schreiber, G.; Fleishman, S. J., Computational design of protein-protein interactions. Curr Opin Struct Biol 2013, 23 (6), 903–10.

3. Ljubetic, A.; Gradisar, H.; Jerala, R., Advances in design of protein folds and assemblies. Curr Opin Chem Biol 2017, 40, 65–71.

4. Lapenta, F.; Aupic, J.; Strmsek, Z.; Jerala, R., Coiled coil protein origami: from modular design principles towards biotechnological applications. Chem Soc Rev 2018, 47 (10), 3530–3542.

5. Voss, S.; Klewer, L.; Wu, Y. W., Chemically induced dimerization: reversible and spatiotemporal control of protein function in cells. Curr Opin Chem Biol 2015, 28, 194–201.

6. Lee, M. J.; Mantell, J.; Brown, I. R.; Fletcher, J. M.; Verkade, P.; Pickersgill, R. W.; Woolfson, D. N.; Frank, S.; Warren, M. J., De novo targeting to the cytoplasmic and luminal side of bacterial microcompartments. Nat Commun 2018, 9 (1), 3413.

7. Lee, M. J.; Mantell, J.; Hodgson, L.; Alibhai, D.; Fletcher, J. M.; Brown, I. R.; Frank, S.; Xue, W. F.; Verkade, P.; Woolfson, D. N.; Warren, M. J., Engineered synthetic scaffolds for organizing proteins within the bacterial cytoplasm. Nat Chem Biol 2018, 14 (2), 142–147.

8. Pu, J.; Zinkus-Boltz, J.; Dickinson, B. C., Evolution of a split RNA polymerase as a versatile biosensor platform. Nat Chem Biol 2017, 13 (4), 432–438.

9. Ptashne, M.; Gann, A., Transcriptional activation by recruitment. Nature 1997, 386 (6625), 569–77.

10. Garvie, C. W.; Wolberger, C., Recognition of specific DNA sequences. Mol Cell 2001, 8 (5), 937–46.

11. Rojo, F., Mechanisms of transcriptional repression. Current Opinion in Microbiology 2001, 4 (2), 145–151.

12. Becker, N. A.; Schwab, T. L.; Clark, K. J.; Maher, L. J., 3rd, Bacterial gene control by DNA looping using engineered dimeric transcription activator like effector (TALE) proteins. Nucleic Acids Res 2018, 46 (5), 2690–2696.

13. Hays, L. B.; Chen, Y. S.; Hu, J. C., Two-hybrid system for characterization of protein-protein interactions in E. coli. Biotechniques 2000, 29 (2), 288–90, 292, 294 passim.

14. Dove, S. L.; Joung, J. K.; Hochschild, A., Activation of prokaryotic transcription through arbitrary protein-protein contacts. Nature 1997, 386 (6625), 627–30.

15. Stynen, B.; Tournu, H.; Tavernier, J.; Van Dijck, P., Diversity in genetic in vivo methods for protein-protein interaction studies: from the yeast two-hybrid system to the mammalian split-luciferase system. Microbiol Mol Biol Rev 2012, 76 (2), 331–82.

16. Hsu, C.; Jaquet, V.; Gencoglu, M.; Becskei, A., Protein Dimerization Generates Bistability in Positive Feedback Loops. Cell reports 2016, 16 (5), 1204–1210.

17. Lupas, A. N.; Bassler, J., Coiled Coils - A Model System for the 21st Century. Trends Biochem Sci 2017, 42 (2), 130–140.

18. Lupas, A. N.; Bassler, J.; Dunin-Horkawicz, S., The Structure and Topology of alpha-Helical Coiled Coils. Subcell Biochem 2017, 82, 95–129.

19. Woolfson, D. N., The design of coiled-coil structures and assemblies. Adv Protein Chem 2005, 70, 79–112.

20. Woolfson, D. N., Coiled-Coil Design: Updated and Upgraded. Subcell Biochem 2017, 82, 35–61.

21. Harbury, P. B.; Zhang, T.; Kim, P. S.; Alber, T., A switch between two-, three-, and four-stranded coiled coils in GCN4 leucine zipper mutants. Science 1993, 262 (5138), 1401–7.

22. Woolfson, D. N.; Bartlett, G. J.; Bruning, M.; Thomson, A. R., New currency for old rope: from coiled-coil assemblies to alpha-helical barrels. Curr Opin Struct Biol 2012, 22 (4), 432–41.

23. Fletcher, J. M.; Boyle, A. L.; Bruning, M.; Bartlett, G. J.; Vincent, T. L.; Zaccai, N. R.; Armstrong, C. T.; Bromley, E. H.; Booth, P. J.; Brady, R. L.; Thomson, A. R.; Woolfson, D. N., A basis set of de novo coiled-coil peptide oligomers for rational protein design and synthetic biology. ACS Synth Biol 2012, 1 (6), 240–50.

24. Thomas, F.; Boyle, A. L.; Burton, A. J.; Woolfson, D. N., A Set of de Novo Designed Parallel Heterodimeric Coiled Coils with Quantified Dissociation Constants in the Micromolar to Sub-nanomolar Regime. J Am Chem Soc 2013, 135, 5161–5166.

25. Thompson, K. E.; Bashor, C. J.; Lim, W. A.; Keating, A. E., SYNZIP Protein Interaction Toolbox: in Vitro and in Vivo Specifications of Heterospecific Coiled-Coil Interaction Domains. ACS Synthetic Biology 2012, 1 (4), 118–129.

26. Negron, C.; Keating, A. E., A set of computationally designed orthogonal antiparallel homodimers that expands the synthetic coiled-coil toolkit. J Am Chem Soc 2014, 136 (47), 16544–56.

27. Aronsson, C.; Danmark, S.; Zhou, F.; Oberg, P.; Enander, K.; Su, H.; Aili, D., Self-sorting heterodimeric coiled coil peptides with defined and tuneable self-assembly properties. Scientific reports 2015, 5, 14063.

28. Hussey, B. J.; McMillen, D. R., Programmable T7-based synthetic transcription factors. Nucleic Acids Res 2018, 46 (18), 9842–9854.

29. Karimova, G.; Pidoux, J.; Ullmann, A.; Ladant, D., A bacterial two-hybrid system based on a reconstituted signal transduction pathway. Proc Natl Acad Sci U S A 1998, 95 (10), 5752–6.

30. Dove, S. L.; Hochschild, A., A bacterial two-hybrid system based on transcription activation. Methods Mol Biol 2004, 261, 231–46.

31. Alberti, S.; Oehler, S.; von Wilcken-Bergmann, B.; Muller-Hill, B., Genetic analysis of the leucine heptad repeats of Lac repressor: evidence for a 4-helical bundle. The EMBO journal 1993, 12 (8), 3227–36.

32. Kercher, M. A.; Lu, P.; Lewis, M., Lac repressor-operator complex. Curr Opin Struct Biol 1997, 7 (1), 76–85.

33. Dong, F.; Spott, S.; Zimmermann, O.; Kisters-Woike, B.; Muller-Hill, B.; Barker, A., Dimerisation mutants of Lac repressor. I. A monomeric mutant, L251A, that binds Lac operator DNA as a dimer. Journal of molecular biology 1999, 290 (3), 653–66.

34. Gaj, T.; Gersbach, C. A.; Barbas, C. F., 3rd, ZFN, TALEN, and CRISPR/Cas-based methods for genome engineering. Trends Biotechnol 2013, 31 (7), 397–405.

35. Politz, M. C.; Copeland, M. F.; Pfleger, B. F., Artificial repressors for controlling gene expression in bacteria. Chemical Communications 2013, 49 (39), 4325–4327.

36. Richter, A.; Streubel, J.; Boch, J., TAL Effector DNA-Binding Principles and Specificity. Methods Mol Biol 2016, 1338, 9–25.

37. Moore, R.; Chandrahas, A.; Bleris, L., Transcription Activator-like Effectors: A Toolkit for Synthetic Biology. ACS Synthetic Biology 2014, 3 (10), 708–716.

38. Khalil, A. S.; Collins, J. J., Synthetic biology: applications come of age. Nat Rev Genet 2010, 11 (5), 367–79.

39. Moon, T. S.; Lou, C.; Tamsir, A.; Stanton, B. C.; Voigt, C. A., Genetic programs constructed from layered logic gates in single cells. Nature 2012.

40. Gardner, T. S.; Cantor, C. R.; Collins, J. J., Construction of a genetic toggle switch in Escherichia coli. Nature 2000, 403 (6767), 339–42.

41. Guet, C. C.; Elowitz, M. B.; Hsing, W.; Leibler, S., Combinatorial synthesis of genetic networks. Science 2002, 296 (5572), 1466–70.

42. Stanton, B. C.; Nielsen, A. A.; Tamsir, A.; Clancy, K.; Peterson, T.; Voigt, C. A., Genomic mining of prokaryotic repressors for orthogonal logic gates. Nat Chem Biol 2014, 10 (2), 99–105.

43. Temme, K.; Hill, R.; Segall-Shapiro, T. H.; Moser, F.; Voigt, C. A., Modular control of multiple pathways using engineered orthogonal T7 polymerases. Nucleic Acids Res 2012, 40 (17), 8773–81.

44. Pellizzoni, M. M.; Schwizer, F.; Wood, C. W.; Sabatino, V.; Cotelle, Y.; Matile, S.; Woolfson, D. N.; Ward, T. R., Chimeric Streptavidins as Host Proteins for Artificial Metalloenzymes. ACS Catalysis 2018, 8 (2), 1476–1484.

45. Wood, C. W.; Heal, J. W.; Thomson, A. R.; Bartlett, G. J.; Ibarra, A. A.; Brady, R. L.; Sessions, R. B.; Woolfson, D. N., ISAMBARD: an open-source computational environment for biomolecular analysis, modelling and design. Bioinformatics 2017, 33 (19), 3043–3050.

46. Battesti, A.; Bouveret, E., The bacterial two-hybrid system based on adenylate cyclase reconstitution in Escherichia coli. Methods 2012, 58 (4), 325–34.

47. Karimova, G.; Ullmann, A.; Ladant, D., Protein-protein interaction between Bacillus stearothermophilus tyrosyl-tRNA synthetase subdomains revealed by a bacterial two-hybrid system. J Mol Microbiol Biotechnol 2001, 3 (1), 73–82.

48. Manelyte, L.; Guy, C. P.; Smith, R. M.; Dillingham, M. S.; McGlynn, P.; Savery, N. J., The unstructured C-terminal extension of UvrD interacts with UvrB, but is dispensable for nucleotide excision repair. DNA repair 2009, 8 (11), 1300–10.

49. Rhodius, V. A.; Segall-Shapiro, T. H.; Sharon, B. D.; Ghodasara, A.; Orlova, E.; Tabakh, H.; Burkhardt, D. H.; Clancy, K.; Peterson, T. C.; Gross, C. A.; Voigt, C. A., Design of orthogonal genetic switches based on a crosstalk map of sigmas, anti-sigmas, and promoters. Mol Syst Biol 2013, 9, 702.

50. Filonov, G. S.; Piatkevich, K. D.; Ting, L. M.; Zhang, J.; Kim, K.; Verkhusha, V. V., Bright and stable near-infrared fluorescent protein for in vivo imaging. Nature biotechnology 2011, 29 (8), 757–61.

51. Thibodeau, S. A.; Fang, R.; Joung, J. K., High-throughput beta-galactosidase assay for bacterial cell-based reporter systems. Biotechniques 2004, 36 (3), 410–5.

52. Bernhardt, T. G.; de Boer, P. A., The Escherichia coli amidase AmiC is a periplasmic septal ring component exported via the twin-arginine transport pathway. Molecular microbiology 2003, 48 (5), 1171–82.

## References

1. Battesti, A.; Bouveret, E., The bacterial two-hybrid system based on adenylate cyclase reconstitution in Escherichia coli. Methods 2012, 58 (4), 325–34.

2. Karimova, G.; Ullmann, A.; Ladant, D., Protein-protein interaction between Bacillus stearothermophilus tyrosyl-tRNA synthetase subdomains revealed by a bacterial two-hybrid system. J Mol Microbiol Biotechnol 2001, 3 (1), 73–82.

3. Manelyte, L.; Guy, C. P.; Smith, R. M.; Dillingham, M. S.; McGlynn, P.; Savery, N. J., The unstructured C-terminal extension of UvrD interacts with UvrB, but is dispensable for nucleotide excision repair. DNA repair 2009, 8 (11), 1300–10.

4. Filonov, G. S.; Piatkevich, K. D.; Ting, L. M.; Zhang, J.; Kim, K.; Verkhusha, V. V., Bright and stable near-infrared fluorescent protein for in vivo imaging. Nature biotechnology 2011, 29 (8), 757–61.

5. Rhodius, V. A.; Segall-Shapiro, T. H.; Sharon, B. D.; Ghodasara, A.; Orlova, E.; Tabakh, H.; Burkhardt, D. H.; Clancy, K.; Peterson, T. C.; Gross, C. A.; Voigt, C. A., Design of orthogonal genetic switches based on a crosstalk map of sigmas, anti-sigmas, and promoters. Mol Syst Biol 2013, 9, 702.

6. Cronan, J. E., A family of arabinose-inducible Escherichia coli expression vectors having pBR322 copy control. Plasmid 2006, 55 (2), 152–157.

7. Sander, J. D.; Cade, L.; Khayter, C.; Reyon, D.; Peterson, R. T.; Joung, J. K.; Yeh, J. R., Targeted gene disruption in somatic zebrafish cells using engineered TALENs. Nature biotechnology 2011, 29 (8), 697–8.

8. Savery, N. J.; Lloyd, G. S.; Kainz, M.; Gaal, T.; Ross, W.; Ebright, R. H.; Gourse, R. L.; Busby, S. J., Transcription activation at Class II CRP-dependent promoters: identification of determinants in the C-terminal domain of the RNA polymerase alpha subunit. EMBO J 1998, 17 (12), 3439–47.

9. Fletcher, J. M.; Boyle, A. L.; Bruning, M.; Bartlett, G. J.; Vincent, T. L.; Zaccai, N. R.; Armstrong, C. T.; Bromley, E. H.; Booth, P. J.; Brady, R. L.; Thomson, A. R.; Woolfson, D. N., A basis set of de novo coiled-coil peptide oligomers for rational protein design and synthetic biology. ACS Synth Biol 2012, 1 (6), 240–50.

10. Thomas, F.; Boyle, A. L.; Burton, A. J.; Woolfson, D. N., A set of de novo designed parallel heterodimeric coiled coils with quantified dissociation constants in the micromolar to sub-nanomolar regime. Journal of the American Chemical Society 2013, 135 (13), 5161–6.

